# Engineered *Methylobacterium extorquens* grows well on methoxylated aromatics due to its formaldehyde metabolism and stress response

**DOI:** 10.1101/2025.03.17.643779

**Authors:** Akorede L Seriki, Alexander B Alleman, Tomislav Ticak, Alyssa C Baugh, Jack W Creagh, Christopher J Marx

## Abstract

Lignin is a vast yet underutilized source of renewable energy. The microbial valorization of lignin is challenging due to the toxicity of its degradation intermediates, particularly formaldehyde. In this study, we engineered *Methylobacterium extorquens* to metabolize lignin-derived methoxylated aromatics, vanillate (VA) and protocatechuate (PCA), by introducing the *van* and *pca* gene clusters. Compared to *Pseudomonas putida*, *M. extorquens* exhibited better formaldehyde detoxification, enabling robust growth on VA without accumulation of formaldehyde. Genetic analyses confirmed that formaldehyde oxidation and stress response systems, rather than C_1_ assimilation, were important for VA metabolism. Additionally, VA and PCA were found to disrupt membrane potential, contributing to their inherent toxicity. Our findings establish *M. extorquens* as a promising chassis for lignin valorization and provide a framework for engineering formaldehyde-resistant microbial platforms.

## Introduction

Lignin, the second most abundant polymer on Earth, is a major structural component of plant biomass and presents a promising source for biofuel production (1, 2). Despite its potential, the complex chemical structure of lignin and the toxicity of its degradation products pose significant challenges to its efficient utilization in biofuel generation (3–5). The biological utilization of lignin is particularly challenging for two main reasons - its structural complexity and resistance to degradation. First, lignin is a highly irregular and heterogeneous polymer, composed of phenylpropanoid units derived from the amino acid phenylalanine. These units, which originate from monolignols (*p*-coumaryl alcohol, coniferyl alcohol, and sinapyl alcohol), differ in their number of hydroxyl groups, oxidation states, and degree of methoxylation. This variability results in the formation of three main lignin subunits: *p*-hydroxyphenyl (H), guaiacyl (G), and syringyl (S) units, making lignin structurally diverse and difficult to process uniformly (6). Second, lignin is polymerized through oxidative radical coupling, a process catalyzed by peroxidases and laccases, forming complex carbon-carbon (C–C) and ether (C–O–C) linkages. This extensive cross-linking contributes to lignin’s rigidity and resistance to enzymatic breakdown. While fungi and some bacteria possess lignin-degrading enzymes—such as laccases, lignin peroxidase, manganese peroxidase, and dye-decolorizing peroxidases—these biological pathways are often slow and incomplete, limiting their efficiency in industrial applications (7–12). Due to these challenges, industrial lignin bioprocessing primarily relies on physicochemical hydrolysis methods, such as acid, alkaline, and oxidative treatments, which more effectively break down lignin (10, 11, 13). However, these chemical processes often produce a complex mixture of aromatic compounds, requiring further refinement for efficient microbial conversion into valuable bioproducts (6, 12, 14).

A variety of organisms can metabolize the diverse cocktail of lignin-derived aromatic compounds. Among these compounds, vanillate (VA) has been widely used as a model compound to study aromatic degradation via the β-ketoadipate pathway (15, 16), which is a well-conserved metabolic route that enables soil bacteria to degrade aromatics. This pathway plays a crucial role in microbial lignin valorization, as it facilitates the conversion of methoxylated and hydroxylated aromatics, such as VA and protocatechuate (PCA), into tricarboxylic acid (TCA) cycle intermediates. The uptake of VA is facilitated by VanK, a specific transporter that enables VA to enter into the cell (17–19). Once inside, VA undergoes *O*-demethylation by vanillate *O*-demethylase (VanAB), producing PCA and releasing formaldehyde, as a toxic metabolic intermediate. (18–20). Meanwhile, PCA serves as a substrate for the β-ketoadipate pathway, undergoing ortho-cleavage by protocatechuate 3,4-dioxygenase, generating β-carboxy-*cis*,*cis*-muconate. This intermediate is subsequently converted into β-ketoadipate *enol*-lactone through a series of enzymatic transformations, primarily involving muconate cycloisomerase and β-ketoadipate *enol*-lactone hydrolase. The resulting β-ketoadipate is further processed by β-ketoadipate succinyl-CoA transferase and β-ketoadipyl-CoA thiolase, ultimately yielding succinyl-CoA and acetyl-CoA, both of which feed into the TCA cycle for energy production and biomass synthesis (21, 22). (Figure 1). The ß-ketoadipate pathway not only allows bacteria to efficiently utilize lignin-derived aromatics as a carbon source but also serves as a crucial link between lignin depolymerization and microbial metabolism. Given its conservation across various bacterial species, the substrates that funnel through the β-ketoadipate pathway are key targets for metabolic engineering aimed at improving microbial lignin valorization for sustainable biochemical production (23). While many bacteria possess the capacity to utilize aromatics, metabolic engineering efforts have focused upon a few model systems, such as *Pseudomonas putida*, as platforms for lignin valorization (24, 25). These efforts have led to an increased breadth of lignin-derived intermediates that can be utilized and a wide variety of bioproducts (26–29).

**Figure 1:**
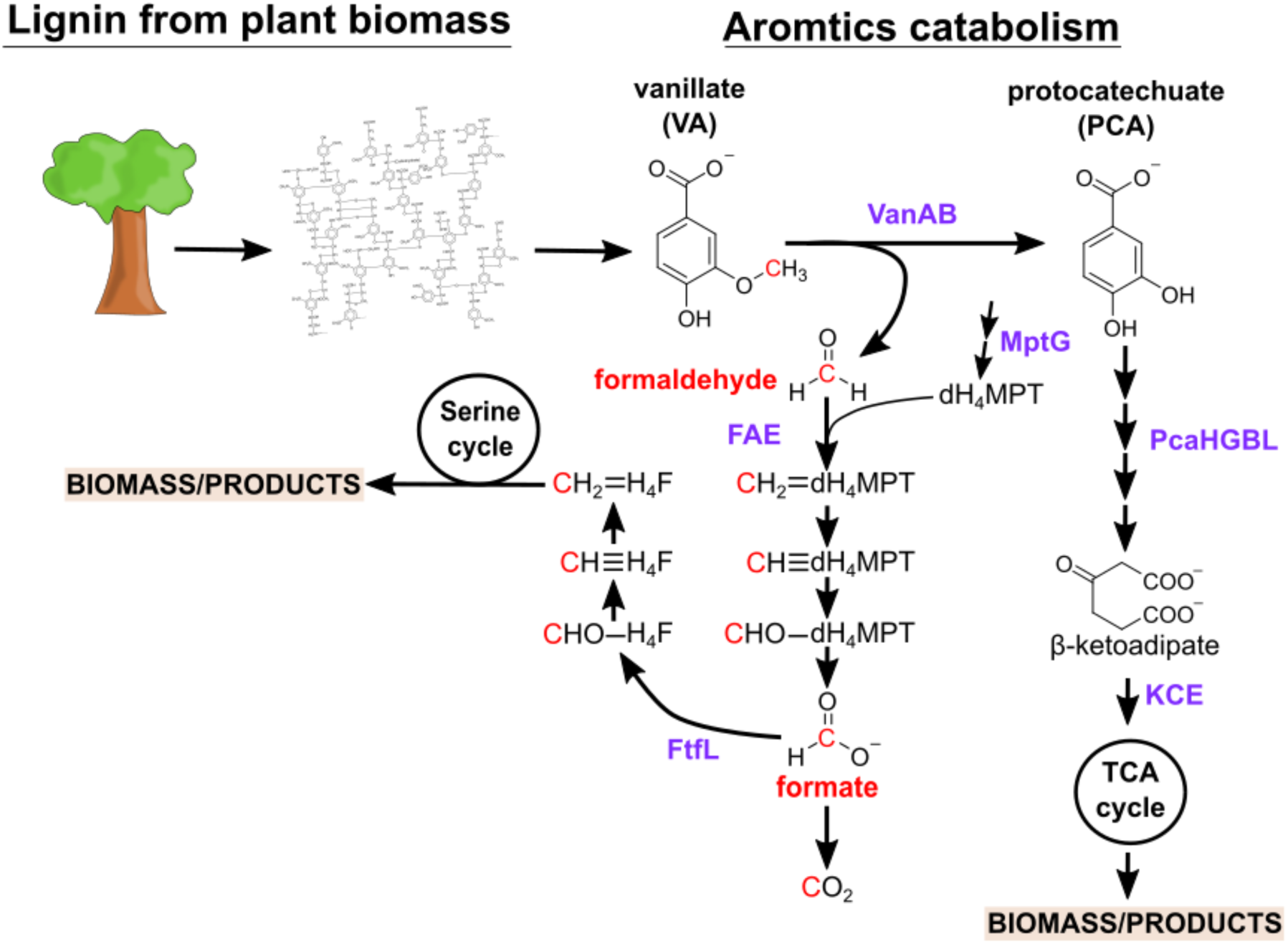
Integrated pathways for the metabolism of lignin-derived aromatics and single-carbon compounds in *Methylobacterium* species. Lignin is depolymerized, releasing a complex mixture of methoxylated aromatic compounds, including vanillate (VA). VA undergoes vanillate demethylation, catalyzed by vanillate monooxygenase (VanAB), converting it into protocatechuate (PCA) and generating formaldehyde – a toxic byproduct. The aromatic rings are cleaved and dissimilated via β-ketoadipate which enters the β-ketoadipate pathway. In *Methylobacterium* species, the usual two-step conversion of β-ketoadipate to TCA cycle intermediates is bypassed by a single-step reaction catalyzed by the homologous enzyme, labelled here as KCE, directly linking β-ketoadipate to the TCA cycle. Meanwhile, formaldehyde is further dissimilated to formate and assimilated via serine cycle into biomass. Key enzymes noted in purple text.

Aromatic compounds, including those derived from lignin, exert toxicity on microbial cells even at low concentrations and this toxicity presents a challenge for efficient bioconversion (30–32). Many of these compounds, such as VA, can disrupt cellular function due to their physicochemical properties. Like other weak organic acids, VA can passively diffuse across bacterial membranes in its protonated form. Once inside the cell, it dissociates due to the higher intracellular pH, leading to cytoplasmic acidification and disruption of the proton motive force, leading to membrane depolarization (33, 34). Additionally, the accumulation of certain aromatic intermediates can impose other metabolic stresses, such as oxidative stress and DNA damage (35, 36), further limiting microbial growth and bioprocessing efficiency.

Another major toxicity challenge with methoxylated aromatics like VA, arises from the fact that their methoxy groups are converted into formaldehyde, which is a highly reactive electrophile that leads to protein and DNA damage (37). Moreso, its accumulation can be lethal if not efficiently removed (38, 39). With methoxy groups being commonly around 15-20% of the total carbon atoms in lignin (40), their breakdown can result in a substantial release of this toxic molecule, posing a major metabolic burden on microbial systems optimized for the utilization of lignin-derived aromatics. For example, despite that fact that *P. putida* can grow on methoxylated versions of lignin-derived compounds (41–44), it has a limited resistance to formaldehyde. Growth of *P. putida* is reduced by 40% at just 0.5 mM formaldehyde (43), and it accumulates formaldehyde during growth on even a low concentration of 2 mM VA (44). This buildup suggests that the formaldehyde detoxification mechanisms of *P. putida* may be insufficient under these conditions, which could pose a challenge for its use in lignin valorization. Given that VA metabolism contributes to accumulation of formaldehyde intracellularly, understanding the interplay of the combined toxicity of VA and formaldehyde is critical for optimizing microbial bioconversion pathways.

Considering the challenge formaldehyde toxicity poses for lignin conversion, it is perhaps worth using a model methylotroph, such as a strain of *Methylobacterium*, as either a conversion chassis or a model system to learn how to improve formaldehyde toxicity in a non-methylotrophic organism. Recently, our lab found that some *Methylobacterium* species grow well on methoxylated aromatic compounds. The presence and organization of the gene clusters enabling growth on methoxylated aromatics vary among *Methylobacterium* strains, being common in some clades of the genus and found only sparsely in other groups (44–46). The gene clusters for aromatics use in *Methylobacterium* contain all the afore-mentioned genes of the canonical β-ketoadipate pathway with the exception of final enzymes, PcaIJ or PcaF. In contrast, the *pca* operon in *Methylobacterium* strains contain a DUF849 predicted to be a β-keto acid cleavage enzyme (which we label as *kce*) (44) (Figure 1). Based upon an earlier study using enzyme screening and active site modeling, this enzyme is proposed to generate succinyl-CoA and acetoacetate from β-ketoadipate and acetyl-CoA (47). The ability to use these compounds is likely because this genus is naturally found on plant surfaces and in soil environments (48), where they would frequently encounter methoxylated aromatics. Critically, we found that, unlike *P. putida*, growth of *Methylobacterium* species on low levels of VA (2 mM) did not lead to formaldehyde accumulation (44). The improved formaldehyde resistance of *M. extorquens* relative to *P. putida* was also observed during growth of a synthetic community on a model mixture of ligno-cellulose monomers (e.g., VA, cellobiose, xylose) where other community members, including *P. putida*, struggled with formaldehyde accumulation (49). Not only could *M. extorquens* grow under these conditions, their presence improved growth of the overall community.

Amongst *Methylobacterium*, formaldehyde metabolism has been best studied in the model organism, *M. extorquens*. This species is used commercially both as a plant-growth promoting microbe for agriculture (50–52) and as a methanol-utilizing organism in bioindustry (53–57). *M. extorquens* oxidizes the formaldehyde produced from C_1_ units such as methanol via the dephospho-tetrahydromethanopterin (dH_4_MPT)-dependent C_1_ carrier pathway to the central branchpoint intermediate, formate (Figure 1) (58, 59). Formate is then either oxidized to CO_2_ to generate ATP and reducing equivalents or assimilated via the tetrahydrofolate (H_4_F)-dependent C_1_ carrier pathway and the serine cycle into biomass (60, 61). Mutants defective in the dH_4_MPT enzymes (e.g., formaldehyde-activating enzyme, encoded by *fae*, (39)), or that lack dH_4_MPT cofactor biosynthesis (e.g., *mptG*, (62, 63)), exhibit extreme sensitivity to methanol due to their inability to efficiently detoxify formaldehyde, leading to its intracellular accumulation and subsequent toxicity (59). In contrast, mutants defective in the H_4_F-dependent pathway (e.g., formate-H_4_F ligase, encoded by *ftfL*, (61)) cannot grow on methanol but are not sensitive to formaldehyde. Collectively, these two pathways contribute to the dynamic ability of the cell to modulate dissimilatory and assimilatory needs without accumulating toxic levels of formaldehyde despite high flux rates (∼2 mM/s) through these pathways (64, 65).

In addition to a dedicated high-capacity formaldehyde oxidation pathway, it has recently been found that *M. extorquens* has a dedicated formaldehyde stress response pathway. Multiple new loci were identified by evolving *M. extorquens* PA1 to increasing formaldehyde concentrations, (66–68). The primary factor implicated was a loss-of-function mutations in Enhanced Formaldehyde Growth protein (encoded by *efgA*), a formaldehyde sensor that directly binds formaldehyde and inhibits translation, leading to growth arrest when intracellular formaldehyde is in excess (68). The role of EfgA with intracellularly-generated formaldehyde appears to be the opposite, whereby its presence aids in switching from succinate to methanol growth, during which time formaldehyde builds up transiently (ref), or in genotypes with mutations in the dH4MPT-dependent formaldehyde oxidation pathway when they are exposed to methanol (ref). Mutations were also observed in EfgB, a putative nucleotide cyclase. The evolved *efgB*^evo1^ allele contributes a slight improvement in formaldehyde resistance on its own, but in combination with the loss-of-function in *efgA*, it greatly enhances formaldehyde resistance (68). It is unclear whether either EfgA or EfgB would play a role in other formaldehyde-producing contexts, such as during growth on methoxylated aromatics.

In order to leverage the knowledge about formaldehyde metabolism in *M. extorquens* PA1 (69–71), this study aimed to introduce the genes for VA metabolism into this background to explore the potential of *M. extorquens* as a model organism for efficiently metabolizing lignin-derived methoxylated aromatics. These genes should also permit growth on the non-methoxylated intermediate, PCA, providing a control as an aromatic that does not generate formaldehyde as an intermediate. We were emboldened by the fact that some closely related natural strains, such as *M. extorquens* AMS-5, are suggested to have acquired aromatic utilization via horizontal gene transfer relatively recently (44).

Remarkably, *M. extorquens* with the *van* and *pca* genes grew at least as well as the organism from which the genes were obtained (*Methylobacterium* nodulans), even on high concentrations of VA. In addition to its robust growth on VA, it also showed substantially greater formaldehyde-resistance than the current model organism, *P. putida*. Our genetic analyses confirm the critical role of the formaldehyde oxidizing dH_4_MPT pathway in VA use and establish that the formaldehyde stress response system is also critical for growth on VA at high concentrations. These insights will be critical to further developing a native methylotroph, like *M. extorquens*, for lignin valorization or may be applied to the formaldehyde resistance of non-methylotrophs such as *P. putida*.

## Methods

### Bacterial strains, media, and chemicals

Strains of *M. extorquens, M. nodulans*, and *P. putida* were all grown in minimal MPIPES medium (67) at 30 °C, whereas *E. coli* was grown in LB at 37 °C. The *M. extorquens* PA1 strains are based upon CM2730, an otherwise wild-type strain that has a Δ*celABC* mutation to remove cellulose biosynthesis and prevent clumping (67). *Methylobacterium* strains were grown on succinate at 3.5 mM, unless otherwise described. For each assay, 10 μL of frozen stock was inoculated into 5 mL MPIPES with 3.5 mM disodium succinate and 50 μg/mL kanamycin (for strains with the pLC291-*pca*-*van* plasmid) and grown 48 h to obtain a stationary phase culture. For growth on aromatics, these stationary phase cells from succinate were diluted into 5 mL MPIPES and either VA or PCA at a low carbon concentration of 5 mM VA or 5.71 mM PCA and grown to stationary phase for a week. Throughout, we use PCA concentrations that are consistently 1/8^th^ higher in concentration than VA to normalize for the number of C atoms per molecule (7 vs. 8, respectively). For growth on the aromatic substrates, cultures were first acclimated to low concentrations of VA (5 mM) or PCA (5.71 mM) to avoid lag from transfer directly from succinate cultures. After reaching the stationary phase, we sub-cultured the cells into sterilized VWR 96-well non-treated, tissue culture plate at 1/64 (v/v) dilution, with each well containing 135 μL fresh medium and the same aromatic substrate as their acclimation but at the tested concentration. For *M. extorquens* with pLC291-*pca-van* we also supplemented the fresh medium with 0.23 ng/mL of anhydrous tetracycline (aTc) to induce a modest level of expression of these genes involved in aromatic compound utilization, as well as 50 μg/mL kanamycin to maintain the plasmid (72).

### Plasmid and strain construction

The plasmid pLC291-*pca-van* was engineered to enable *Methylobacterium extorquens* to metabolize VA or PCA. The backbone vector, pLC291, is an aTc-inducible expression vector previously described by Chubiz et al. (72) that is based upon a small IncP replicon (73). Genes involved in VA and PCA degradation were PCR-amplified from genomic DNA isolated from *M. nodulans* ORS2060. The following genes were amplified and cloned via Gibson assembly into the linearized pLC291 vector downstream of the *P_R_/tetO* promoter region: *vanA*, *vanB*, and *vanK*, which are essential for conversion of VA to PCA (19), and the PCA utilization cluster, corresponding to *pcaHGBD* genes and the *kce* homolog. The Gibson assembly mixtures were transformed into *E. coli* DH5α cells, and transformants were selected on LB agar plates containing 50 μg/mL kanamycin. Colony PCR and restriction digestion analysis were used for initial screening, and plasmid DNA from confirmed clones was sequenced to verify. The complete sequence of pLC291-*pca-van* has been deposited on NCBI (#XXXX). The verified plasmid was introduced into various strains of *M. extorquens* PA1 via triparental conjugation using the helper plasmid pRK2073 (74). Transconjugants were selected on MPIPES succinate plates containing 50 μg/mL kanamycin and verified by Sanger sequencing.

**Table 1:** Strains used in this study.

| Strain | Relevant characteristics | Source or reference |
| --- | --- | --- |
| CM2730 | ‘wild-type’ $\Delta celABC$ *M. extorquens* PA1 | (67) |
| CM3745 | $\Delta efgA$ in CM2730 | (68) |
| CM3753 | $\Delta fae$ in CM2730 | (39) |
| CM3773 | $\Delta ftfl$ in CM2730 | (61) |
| CM3803 | $\Delta mptG$ in CM2730 | (63) |
| CM4669 | CM2730 with pLC291-*pca-van* | This study |
| CM4670 | CM3745 with pLC291-*pca-van* | This study |
| CM4730 | $\Delta efgA$ *efgB*<sup>evol</sup> in CM2730 | (68) |
| CM4770 | *P. putida* KT2440 | (75) |
| CM4913 | NCM3722 *E. coli* K-12 | (76) |
| CM4925 | *M. nodulans* ORS 2060 | (77) |
| CM5110 | CM3753 with pLC291-*pca-van* | This study |
| CM5111 | CM3773 with pLC291-*pca-van* | This study |
| CM5114 | CM3803 with pLC291-*pca-van* | This study |
| CM5118 | CM4730 with pLC291-*pca-van* | This study |

### Growth quantification and measurement of specific growth rate

Optical density (OD) measurements were recorded in a 96-well plate using a Synergy H1 Plate reader (BioTek), at 600 nm (OD_600_). The protocol was set to double orbital continuous shaking at 548 cpm in fast mode, with OD_600_ measurements taken at 15 min intervals, to monitor bacterial growth on aromatics. Raw OD values were processed using R (version 2023.06.1+524) with the growthrates, tidyverse, and ggplot2 packages. Blank wells, containing only media and carbon source without cells, were used to correct for background absorbance. The blank OD value was subtracted from all sample readings to ensure accurate growth estimations. Growth rate estimation was performed by fitting OD_600_ readings from the log-linear growth phase with an exponential growth model in the growthrates package. Growth rates reported for each strain and condition represent the mean of three technical replicates.

### Formaldehyde quantification

Formaldehyde stock (1 M) solution was prepared fresh daily by combining 0.3 g of paraformaldehyde powder (Sigma Aldrich), 9.95 mL of ultrapure water, and 50 μL of 10 N NaOH solution in a sealed tube. The solution was then immersed in a boiling water bath for 20 min to induce depolymerization. Formaldehyde concentration was measured using the method described by Nash (78). In brief, equal volumes of each sample and Nash Reagent B (2 M ammonium acetate, 50 mM glacial acetic acid, 20 mM acetylacetone) were combined in a 96-well plate and incubated at 60 °C for 10 min. Absorbance was measured at 432 nm on a Synergy H1 Plate reader (BioTek). Each plate included samples in triplicate along with a standard curve, also run in triplicate.

### Membrane depolarization assay

Membrane potential was determined using the BacLight^TM^ Bacterial Membrane Potential Kit (Invitrogen). Briefly, the fluorescent probe 3,3’-diethyloxacarbocyanine iodide (DiOC_2_(3)) was applied to determine the changes of bacterial membrane potential. Bacteria in the logarithmic phase of growth were harvested by centrifugation at 4000 ×g for 10 min, and the bacterial concentration was adjusted to 1 × 10^5^ CFU/mL, by diluting in PBS. The bacterial suspensions were incubated with VA/PCA, and appropriate controls, for 30 min and then incubated with DiOC_2_(3) at 25 °C before measuring fluorescence via flow cytometry (Beckman Coulter Cytoflex S).

## Results

### Introduction of the *pca* and *van* genes into *Methylobacterium extorquens* enables growth on vanillate (VA) and protocatechuate (PCA)

Analogous to natural examples of *M. extorquens* strains that appear to have received aromatic utilization clusters via HGT (44), we hypothesized that we could successfully move the *van-pca* gene cluster from *M. nodulans* ORS 2060 (77) into the model strain, *M. extorquens* PA1 (67). Two segments of the *M. nodulans* genome encoding *vanABK* and *pcaHGBL* with a possible *kce* gene were cloned into the regulated expression vector, pLC291 (72). The resulting plasmid, pLC291-*pca-van*, allows for aTc regulated expression of each of these operons under a separate inducible *P_R_/tetO* promoter (Figure 2). Introduction of pLC291-*pca-van* into *M. extorquens* (hereafter referred to as the engineered *M. extorquens*) enabled it to grow on both VA and PCA (Figure 3). While growth on PCA was substantially slower than that of *M. nodulans,* growth on VA at a higher concentration (40 mM) was equivalent and did not exhibit the clumping observed for *M. nodulans*. In the following work, we focus on determining the role of substrate concentration, methylotrophic metabolism, and formaldehyde stress response upon the ability of *M. extorquens* to metabolize methoxylated aromatics.

**Figure 2:**
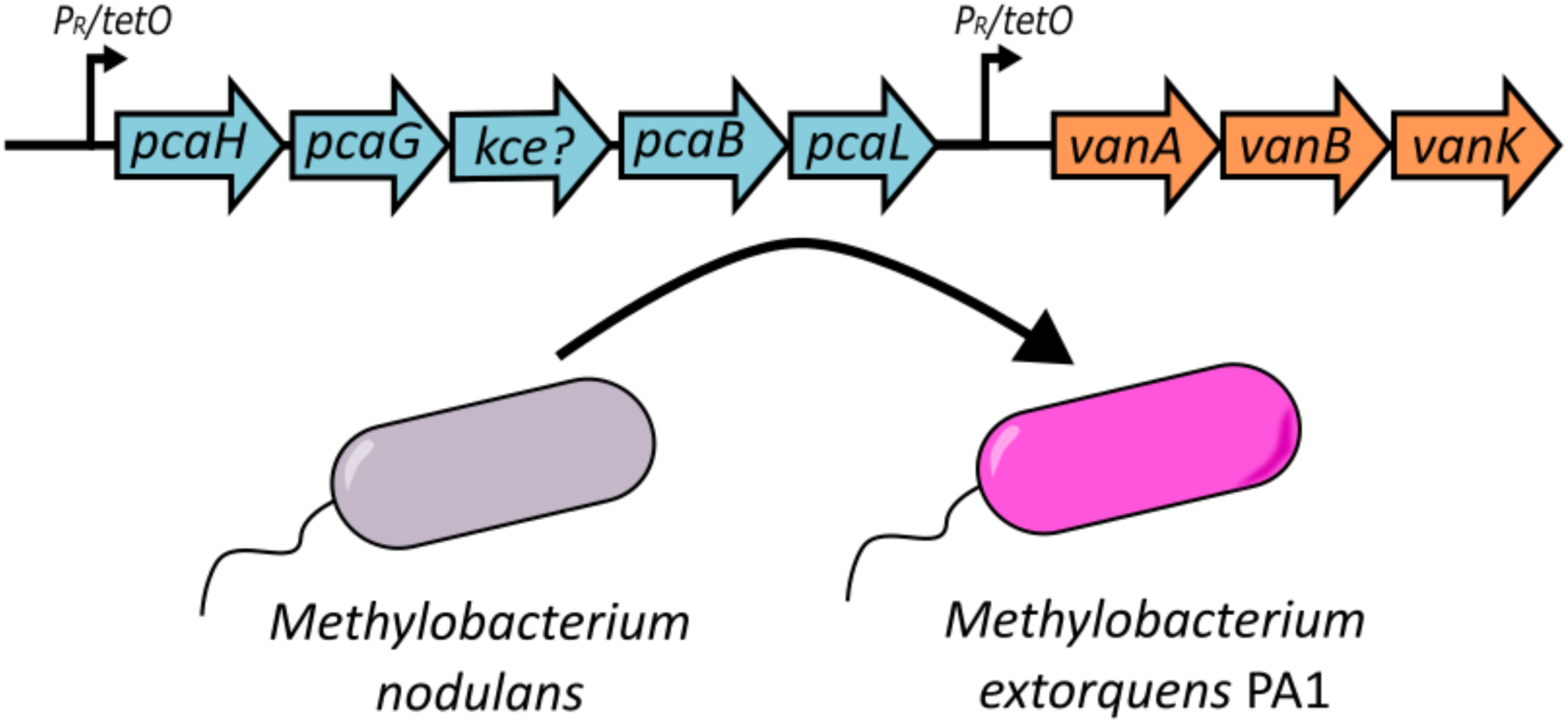
Genetic organization of the pLC291-*pca-van* construct in *Methylobacterium extorquens*. Schematic representation of the cloned gene cluster in pLC291-*pca-van*, showing the *pcaHGBD*+*kce* genes involved in PCA degradation and the *vanABK* genes responsible for VA metabolism. These genes were obtained from *M. nodulans* ORS 2060 and introduced into *M. extorquens* PA1 under the control of the *P_R_/tetO* inducible promoter system.

**Figure 3:**
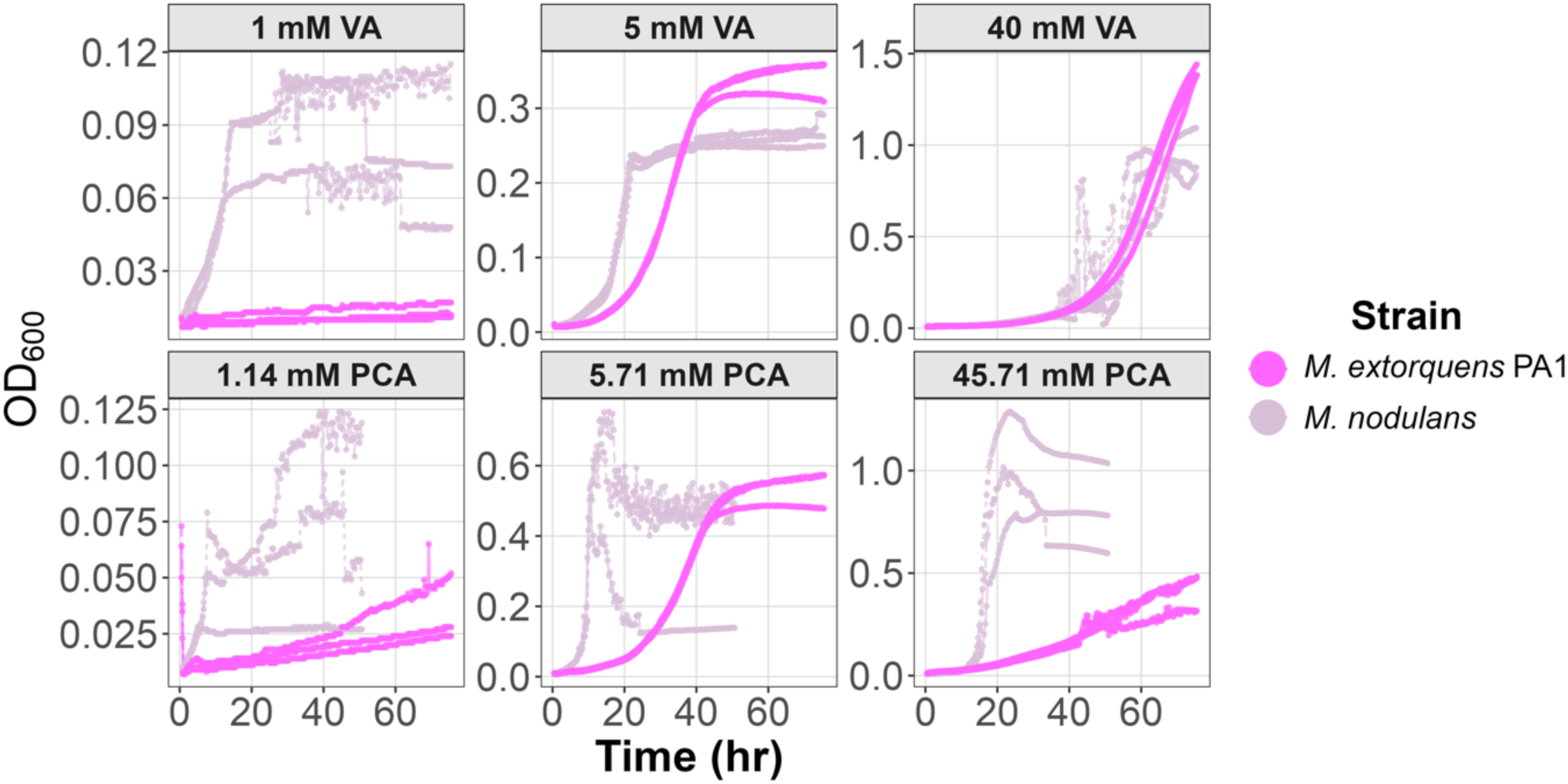
Growth of *M extorquens* carrying pLC291-*pca-van* and *M. nodulans* on low to high concentrations of VA and PCA. *M. nodulans* exhibited faster growth on PCA but showed significant clumping, leading to irregular OD₆₀₀ readings. In contrast, engineered *M. extorquens* maintained a smoother growth curve across all conditions. At high VA concentrations (40 mM), both strains displayed similar growth profiles, but *M. nodulans* again exhibited clumping. The activity of the transporter *VanK* is evident in *M. nodulans* as shown by the growth observed at low concentration (1 mM VA). Three replicates of each strain are shown. The full range of concentrations tested are shown in Figure S1.

### Concentration-dependent growth rate for engineered *M. extorquens* suggests transport via diffusion

To explore the potential of engineered *Methylobacterium extorquens* for bioconversion applications, we investigated how the engineered strain responds to varying concentrations of methoxylated aromatic compounds. We examined growth on succinate, a C_4_-dicarboxylate that does not generate formaldehyde as an intermediate and has a known transporter (79), as a control for what to expect in the absence of toxicity and the presence of active transport with a low Kₘ. As expected, growth rate remained steady across a range of concentrations from 0.5 to 40 mM, with an average rate of approximately 0.20 hr⁻¹ (Figure 4A).

**Figure 4:**
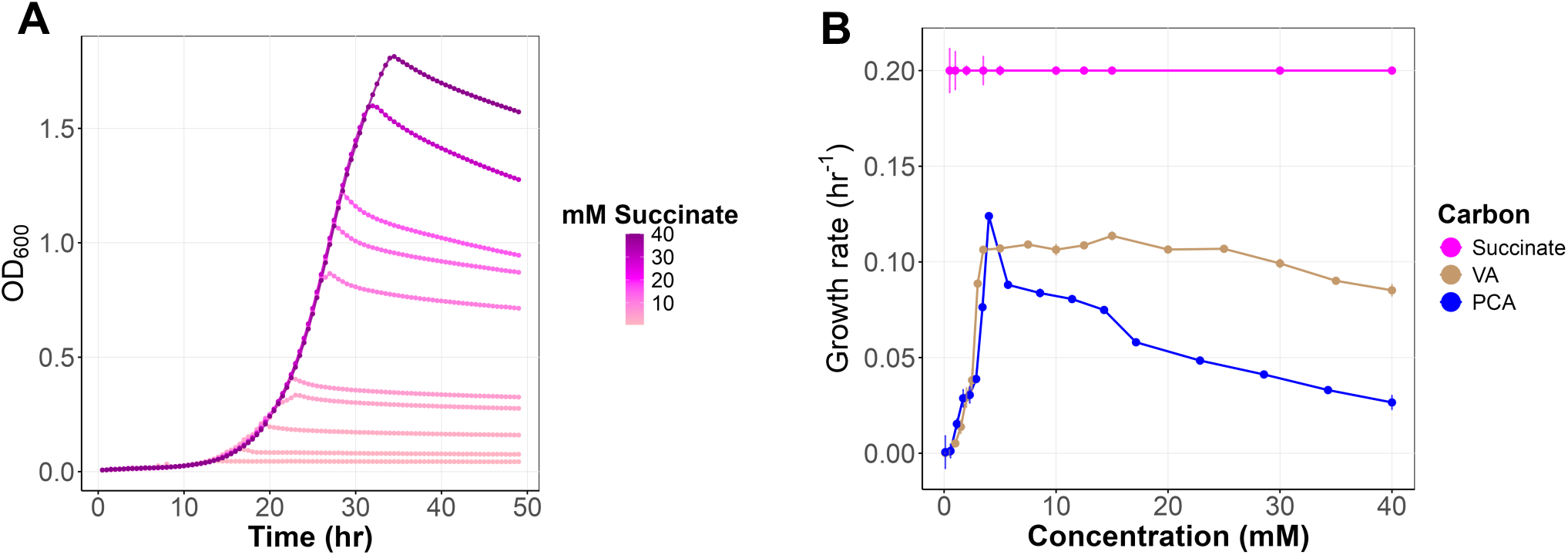
Growth curves of *M. extorquens* across a range of succinate concentrations and growth rates at varying concentrations of succinate, VA, and PCA. **(A)** Growth curves on succinate across a concentration range of 0 to 40 mM, exhibiting stable growth independent of concentration. Data are averages of 3 replicates. (B) Average growth rates (hr⁻¹) of *M. extorquens* on PCA, VA, and succinate as a function of concentration. PCA (blue) and VA (brown) show concentration-dependent inhibition, while succinate (magenta) demonstrates concentration-independent growth rate. Data are averages of three replicates and error bars denote 95% confidence intervals.

In contrast to succinate, the growth of engineered *M. extorquens* using VA and PCA as substrates was variable across concentrations. Very slow growth was seen at low concentrations, and the growth rate increased approximately linearly with either VA or PCA, reaching a peak at 5 mM for either substrate. The linear relationship between growth rate and substrate concentration, and the high substrate concentration for half-maximal growth are inconsistent with active transport being the primary means of uptake. This instead suggests that these substrates are driven into the cell via passive diffusion. PCA appears to exhibit toxic effects beginning at modest concentrations, as growth rate drops sharply past ∼5 mM, with an approximately linear decline in growth rate down to 40% of the maximal rate by 40 mM (Figure 4B). After an initial linear increase with concentration, the growth rate on VA was consistent up to 25 mM before a linear decline at higher concentrations (Figure 4A). Such a concentration-dependent inhibition suggests passive diffusion at lower concentrations, followed by intracellular accumulation to toxic levels at high concentrations, rather than a saturable transport process with a defined affinity (Kₘ).

### *P. putida* accumulates formaldehyde during growth, with increasing concentrations of VA, unlike *M. extorquens*

Discovering that the engineered strain grew on high concentrations of VA (>5 mM), we sought to determine whether there was formaldehyde accumulation at this level. As described above, natural *Methylobacterium* strains were found to avoid formaldehyde accumulation compared to *P. putida*, (but this was only tested at a relatively low concentration of 2 mM (44)). Consistent with what was found for natural *Methylobacterium* strains, our engineered strain grew on 5 mM VA with no formaldehyde accumulation. We then tested growth on higher concentrations of VA and found no formaldehyde accumulation even at 20 mM (Figure 5). We observed similar dynamics at 5 mM VA and at VA concentrations of 20 mM *P. putida* not only failed to grow in duration of exposure (30 hr) but accumulated even higher levels of formaldehyde (Figure 5). This indicates a substantial increase in per capita formaldehyde accumulation which is the likely cause of the inability of *P. putida* to grow at high VA concentrations. These results highlight that, even without any further development, our engineered *M.* e*xtorquens* is superior at dealing with formaldehyde toxicity compared to *P. putida*, and capable of growth at high concentrations of VA.

**Figure 5:**
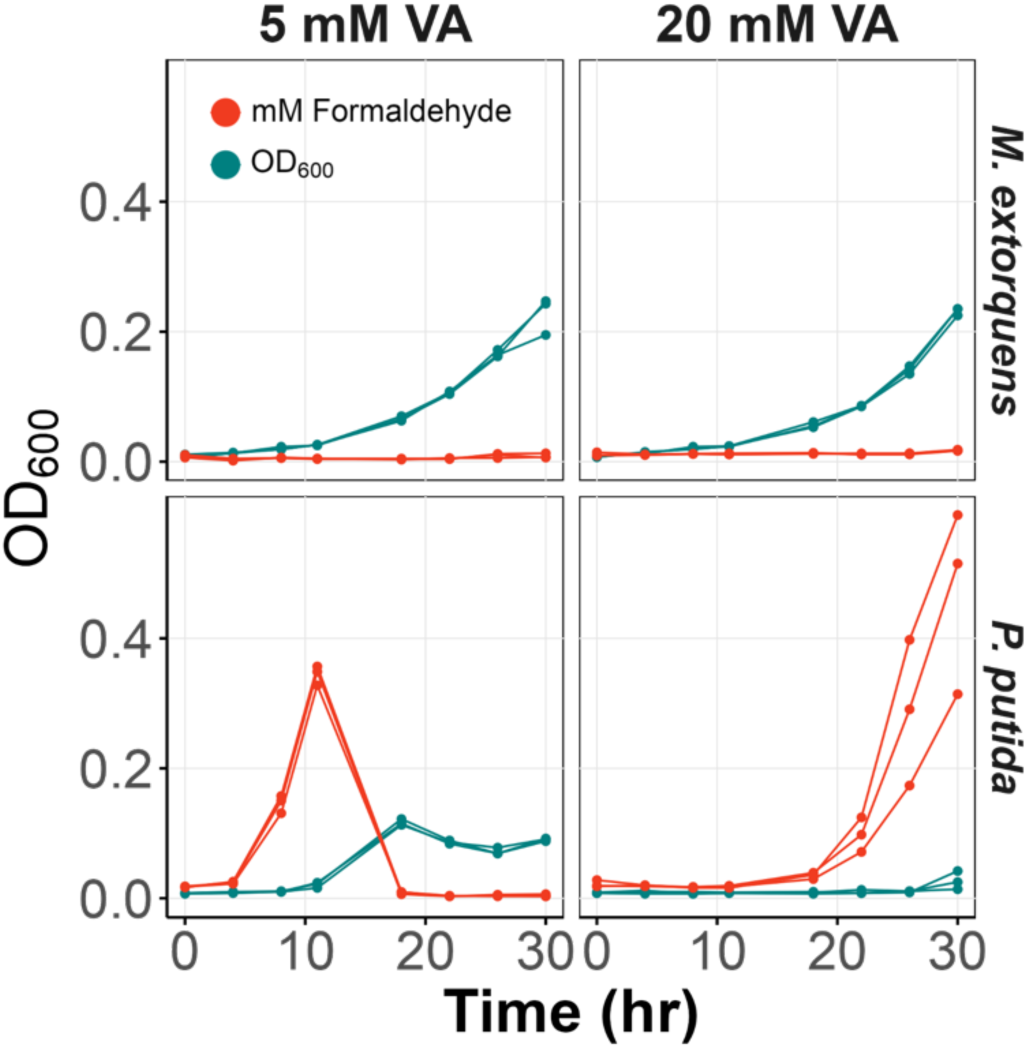
**Formaldehyde accumulation and growth on VA by *M. extorquens* and *P. putida***. Growth (teal) and extracellular formaldehyde levels (red) of *M. extorquens* and *P. putida* cultured in MPIPES supplemented with 5 mM or 20 mM vanillate (VA). *M. extorquens* exhibited no formaldehyde accumulation while *P. putida* accumulated substantial formaldehyde, particularly at 20 mM VA, leading to growth inhibition. Three replicates of each strain are shown.

### Formaldehyde oxidation, but not C₁ assimilation, is essential for growth on methoxylated aromatics in *Methylobacterium extorquens*

To understand the capacity of engineered *M. extorquens* to grow on high concentrations of VA without accumulating formaldehyde, we examined the role of genes central for C_1_ dissimilation and assimilation. To elucidate the role of C_1_ dissimilation, we tested strains with deletions of either Fae or MptG, i.e., the dH₄MPT-dependent pathway for formaldehyde oxidation. For the role of C_1_ assimilation, we tested strains lacking FtfL, which blocks assimilation of formate, by the H_4_F-dependent pathway and serine cycle, into biomass (Figure 1). Mutants in dissimilation and assimilation had remarkably different phenotypes.

The formaldehyde oxidation mutants (Δi*fae* and Δi*mptG*) were unable to grow on VA at any concentration (Figure 6A). The formaldehyde oxidation mutants were able to grow on PCA (Figure S2), confirming that the inability to grow on VA was due to formaldehyde production from VA.

**Figure 6:**
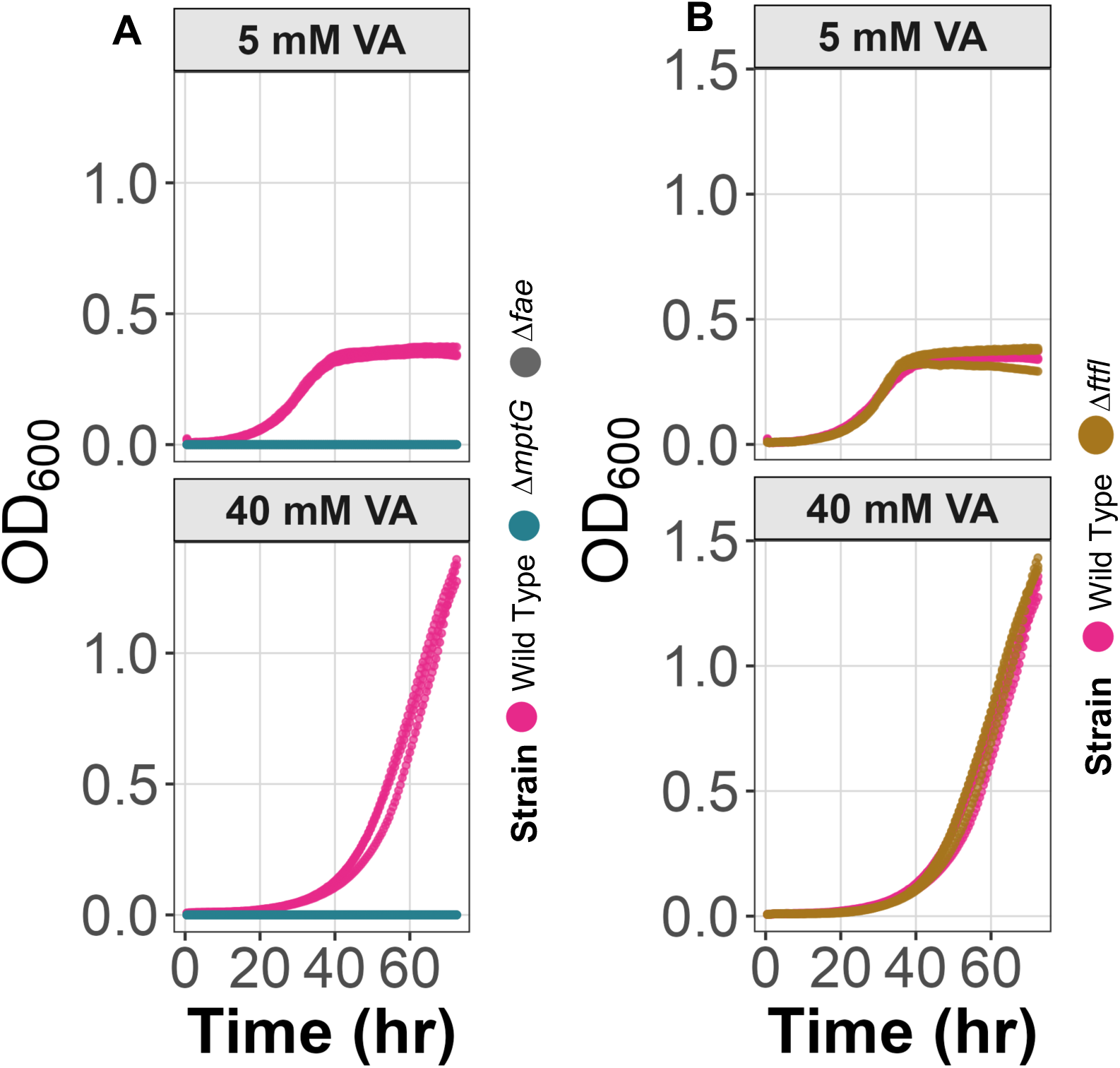
Formaldehyde oxidation but not assimilation contributes to growth of engineered *M. extorquens* on VA. (A) Mutant strains defective in formaldehyde oxidation pathways (Δ*fae,* Δ*mptG*) show impaired growth on VA at both low and high concentrations, highlighting the importance of formaldehyde detoxification in the metabolism of methoxylated aromatics. (B) In contrast, mutants defective in C₁ assimilation pathway (Δ*ftfL*) demonstrate no growth defects on VA, indicating that formate produced from methoxy groups is primarily dissimilated. Three replicates of each strain are shown.

The Δ*ftfL* strain (defective assimilatory module) exhibited a growth profile similar to the engineered strain on VA and even showed a slight growth advantage on PCA (Figures 6B, S2). The similar profile to engineered strain on VA indicates that assimilation of the methoxy group is not essential for growth on VA.

### Mutations beneficial for growth on formaldehyde impart a growth advantage on high concentrations of VA

Given that formaldehyde oxidation is specifically essential to growth on VA compared to PCA, we extended our genetic analysis to consider recently discovered formaldehyde stress response pathways (68–70). We introduced the pLC291-*pca-van* plasmid into backgrounds containing beneficial mutations acquired during evolution on formaldehyde where these pathways were discovered, with the expectation that it will enhance growth on VA but not PCA. Furthermore, we hypothesized that the extent of this benefit should scale with VA concentration. Consistent with these hypotheses, the Δ*efgA* mutation and the Δ*efgA efgB^evo1^* combination conferred an advantage on VA, with larger benefits seen at higher VA concentrations (Figure 7A). In contrast, these mutations had no consistent effect on PCA (Figure S2). The magnitude of advantage on high VA scaled with advantage seen during formaldehyde growth, such as the Δ*efgA efgB^evo1^* strain with two beneficial mutations outperforming the Δ*efgA* mutant (Figure 7B). Taken together, these results highlight that the advantage of *M. extorquens* to grow on VA compared to *P. putida* is due to both its metabolic pathways and formaldehyde stress response systems.

**Figure 7:**
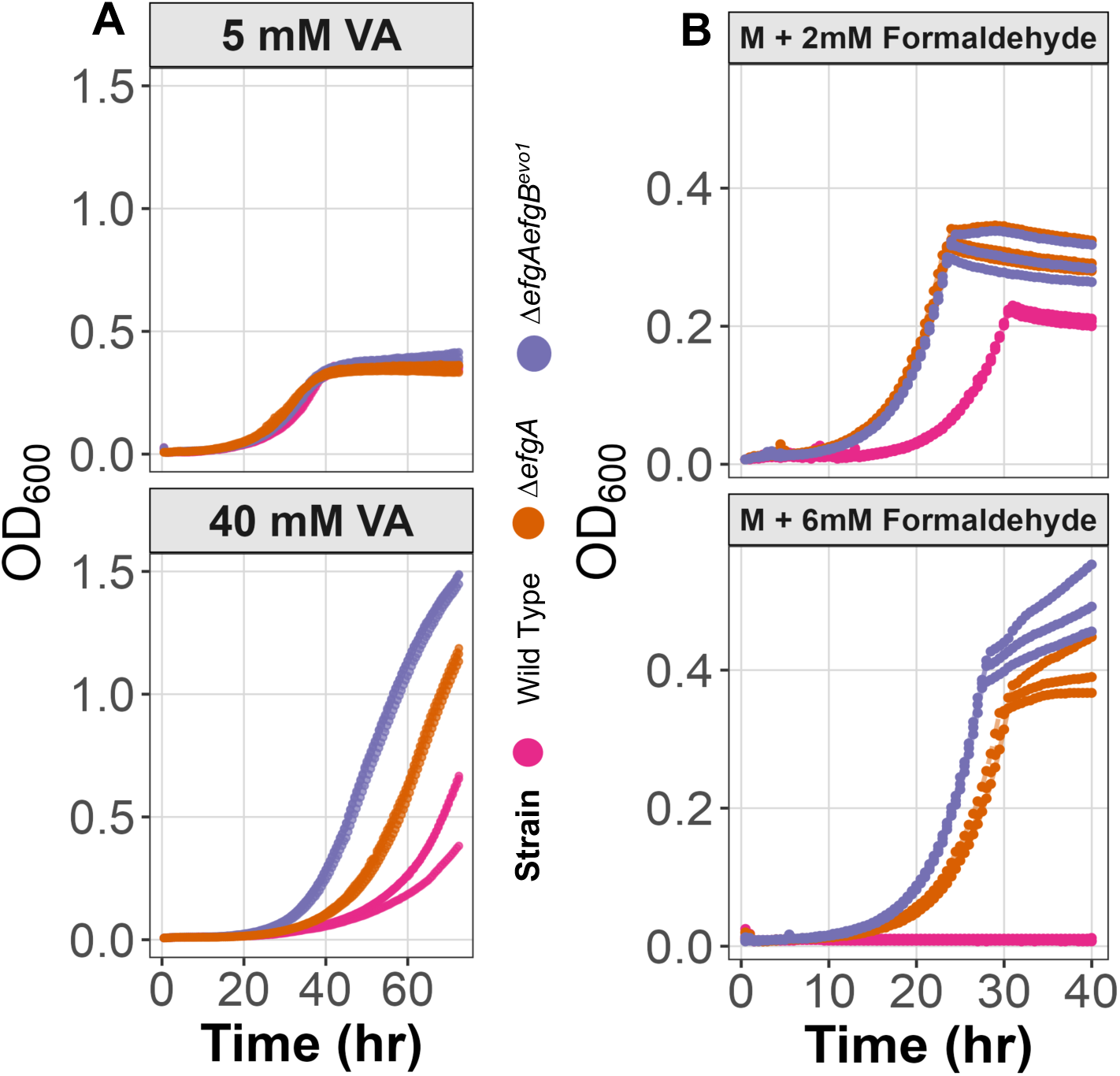
Strains with formaldehyde-evolved alleles of *efgA* or *efgB* exhibit improved growth on VA but not on PCA. Growth curves (OD_600_) of *Methylobacterium extorquens* strains cultured on VA (A) or methanol (M) supplemented with formaldehyde (B) at varying concentrations. Wild-type, Δ*efgA,* and Δ*efgA efgB^evo1^* double mutant were tested across different conditions. The Δ*efgA* mutation conferred a growth advantage at high VA concentrations, and the Δ*efgA efgB^evo1^*double mutant displayed the most significant improvement under both high VA and formaldehyde stress conditions. Three replicates of each strain are shown.

### Aromatic substrate toxicity in non-engineered *M. extorquens*

In addition to formaldehyde toxicity, aromatics such as VA and PCA can also be toxic due to their inherent chemical properties. To determine whether *M. extorquens* is inherently well-suited to withstanding aromatics toxicity, we examined toxicity in wild-type (WT) *M. extorquens* (i.e., lacking pLC291-*pca-van* and therefore unable to metabolize VA and PCA) while growing on succinate, supplemented with VA or PCA. We hypothesize that if toxicity were due to conversion, then inhibition of WT would be lessened, or eliminated. In contrast, if the compounds were toxic whether or not they were metabolized, the strength of inhibition would be similarly strong. To our surprise, both VA and PCA were substantially more toxic to the WT lacking the *pca-van* pathways (Figure 8A). Unlike the engineered strain, growth rate decreased sharply and linearly with increasing VA concentrations (Figure 8B). Similarly, when the wild-type strain was grown on 3.5 mM succinate with increasing PCA concentrations, growth inhibition was also concentration-dependent (Figures 8C and D). These findings indicate that VA and PCA impose direct toxicity beyond formaldehyde accumulation, suggesting additional inhibitory effects on growth. Moreover, this toxicity is exacerbated in the absence of a functional degradation pathway, significantly impairing growth when VA and PCA cannot be metabolized.

**Figure 8:**
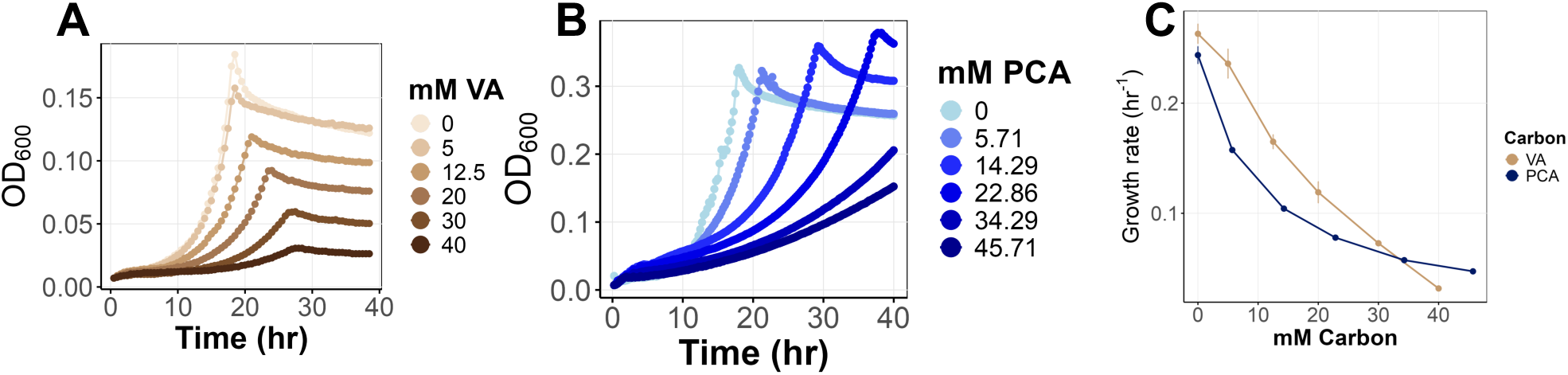
Non-engineered wild type *M. extorquens* is highly sensitive to either VA or PCA during growth on succinate. (A and B) Growth curves and rate of engineered *M. extorquens* on VA with 3.5 mM succinate as a co-substrate, across VA concentrations from 0 to 40 mM, showing severe growth inhibition at higher concentrations. (C) Growth rates of engineered *M. extorquens* on PCA with 3.5 mM succinate as a co-substrate, across PCA concentrations from 0 to 45.71 mM, highlighting the inhibitory effects of increasing PCA concentration. Data are averages of three replicates and error bars denote 95% confidence intervals.

### VA and PCA disrupt membrane potential in *Escherichia coli*

One possible mechanism of aromatic acid toxicity is due to the impact on membrane potential of aromatic acids entering the cell in their protonated form and diffusing out with counterions such as Na^+^ or K^+^. This leads to ion imbalance and dissipation of the proton motive force, ultimately impairing cellular energy production and homeostasis. To assess whether VA or PCA have the ability to impact membrane potential in this manner, we used 3,3’-diethyloxacarbocyanine iodide (DiOC₂(3)) as a membrane potential-sensitive dye that exhibits a red-to-green fluorescence shift in response to changes in membrane polarization. A higher red/green fluorescence ratio indicates a more negative membrane potential (higher ΔΨ), while a lower ratio reflects membrane depolarization. Repeated attempts to apply this dye with *M. extorquens* were unsuccessful, perhaps due to poor diffusion into the cell. As a proxy, we used *Escherichia coli*, since DiOC_2_ has been used extensively to study membrane potential in *E. coli* (80, 81). The assay was first validated using carbonyl cyanide *m*-chlorophenyl hydrazone (CCCP), a known protonophore which depolarizes membranes. CCCP treatment decreased the red/green ratio compared to the untreated control, which was used as a baseline (Figure 9), confirming the dye’s effectiveness in detecting membrane potential changes. Exposure to high concentrations of VA or PCA resulted in a reduction in the red/green fluorescence ratio, comparable to the CCCP-treated condition, indicating strong membrane depolarization. The reduction was moderate at lower concentrations of either aromatic molecule, indicating a dose-dependent effect (Figure 9). These findings align with growth inhibition data, supporting the idea that toxicity at high aromatic concentrations is linked to disruption of membrane potential.

**Figure 9:**
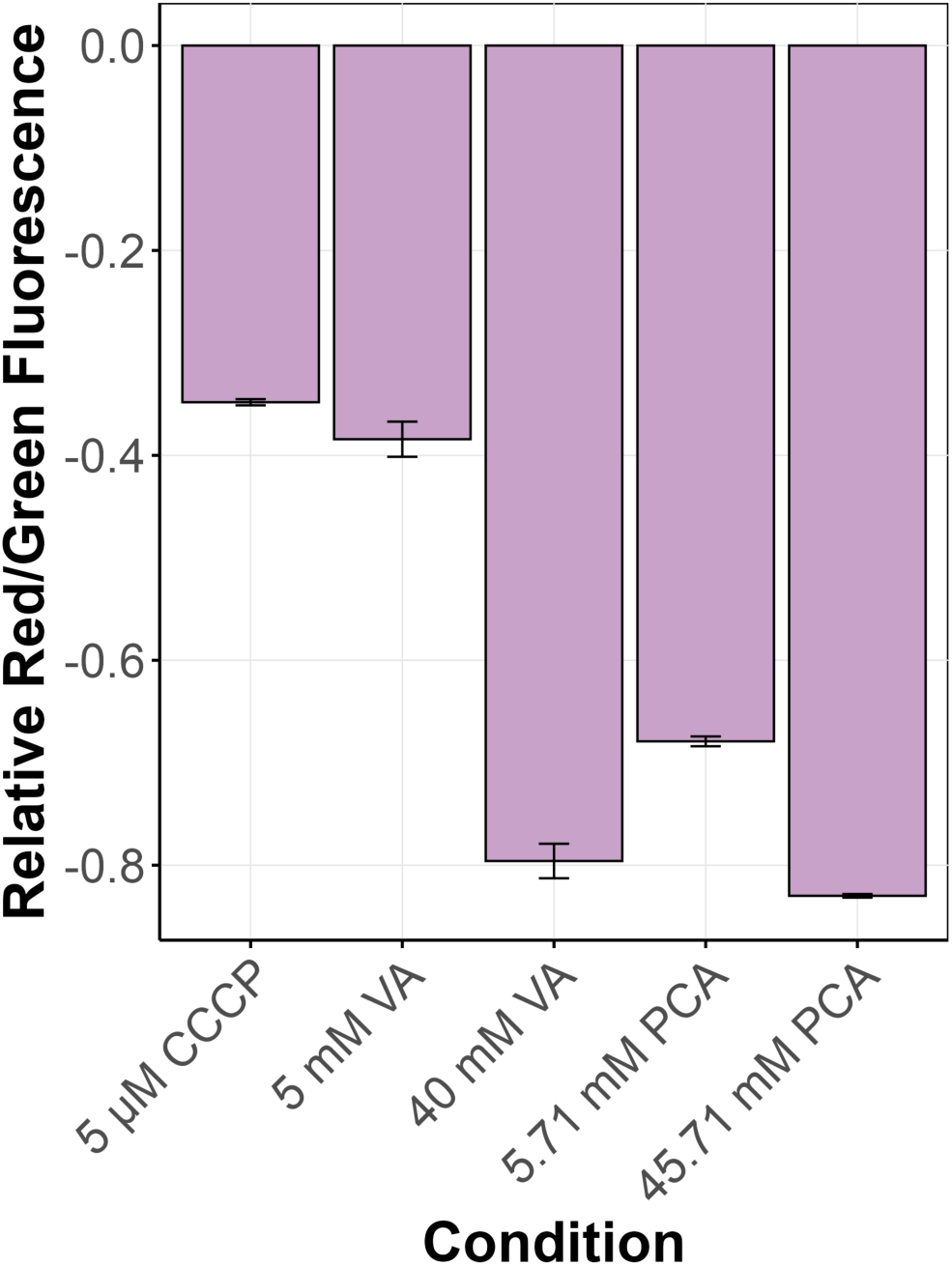
Membrane depolarization in *E. coli* exposed to VA or PCA. Red/green fluorescence ratio of DiOC_2_(3) in *E. coli* cells treated with 5 μM CCCP (positive control for depolarization), VA and PCA. Fluorescence is normalized to DiOC_2_(3) stained cells with no CCCP. Lower (more negative) fluorescence values indicate greater membrane depolarization. CCCP disrupted membrane potential equivalent to a low concentration of VA, whereas higher VA or PCA induced even more depolarization, suggesting that aromatic compounds impact *E. coli* membrane potential in a concentration-dependent manner. Bar graphs represent the average of three replicates and error bars denote 95% confidence intervals.

## Discussion

Our study indicates that introducing the genes for metabolism of methoxylated aromatics into *M. extorquens* PA1 generates a new microbial platform for industrial lignin valorization by avoiding traditional formaldehyde toxicity. Even without additional genetic modifications, this strain grows on high VA concentrations as well as naturally aromatic-utilizing *Methylobacterium* species, such as *M. nodulans*. Furthermore, *M. extorquens* with pLC291-*pca-van* can specifically handle methoxylated aromatics far better than *P. putida*. This positions *M. extorquens* as a promising candidate for lignin valorization and bioremediation applications and suggests that learning what functions underlie its ability to handle high VA could be useful in developing other non-methylotrophic hosts to resist formaldehyde toxicity.

Multiple aspects of our data suggest that transport may be a critical limitation to growth of *M. extorquens* with pLC291-*pca-van* on VA. Unlike growth on succinate, which has a growth rate independent of concentration across the wide range of 0.5-40 mM, both VA and PCA showed a very different pattern. The linear increase in growth up to 5 mM is rather inconsistent with a saturatable, active transporter and suggests it enters the cell via passive diffusion. Given the established capacity of aromatic acids to cross membranes (82), this is unsurprising. This also suggests that, despite the expectation that the VanK transporter should be expressed by the introduced *vanABK* operon, this transporter is either inactive or under-expressed and could be a target for future strain development.

Genetic analyses of mutants defective for formaldehyde dissimilation or assimilation indicated very clearly that formaldehyde oxidation is essential for VA use, but that the released methoxy group does not need to be assimilated into biomass. On the contrary, deleting *ftfL* was beneficial on PCA, which is similar to previous observations in *M. extorquens* AM1 that evolved on succinate and acquired loss-of-function mutations in *ftfL* that benefitted growth on succinate (83). Additionally, growth on VA can be considered similar to growth on a mixture of succinate and methanol, as VA catabolism generates TCA cycle and C_1_ intermediates. *M. extorquens* has been shown to grow faster on a mixture of succinate and methanol than on either substrate alone. Under these conditions, it shifts its metabolism to primarily dissimilate C_1_ units from methanol while increasing the proportion of succinate allocated to biomass production compared to growth on succinate alone (84).

Analysis of engineered *M. extorquens* strains carrying beneficial mutations from formaldehyde stress evolution revealed that the same alleles were beneficial at high VA concentrations. This provides the first linkage of the formaldehyde stress response to multi-C substrates that also contain C_1_ units. Given that there are many methoxylated compounds in the environment, as well as methyl groups found on N or S atoms that are similarly available to convert into formaldehyde, these stress response systems may be broadly important in nature.

The extreme sensitivity of wild-type *M. extorquens* lacking any aromatic metabolic pathways to either VA or PCA was rather surprising. Metabolic conversion appears to not only enable growth but also provide a resistance mechanism. In the absence of further metabolic conversion, VA and PCA were fairly similar in their toxicity. This could be due to a direct effect of either aromatic acid, or it could be due to their known ability to depolarize membranes by escaping from cells with a counter ion. Given that the first step of PCA metabolism is ortho-cleavage of the ring, this would eliminate the aromaticity of the molecule and instead generate a tricarboxylic acid (β-carboxy-*cis*,*cis*-muconate) that would be much less membrane permeable. Using *E. coli* as a proxy, we were able to confirm that VA and PCA are both potent agents for membrane depolarization, each generating stronger effects than the CCCP control even at low (5/5.71 mM) concentrations.

Relative to other foreign metabolic functions introduced into *M. extorquens* (85), the pathway for VA and PCA growth was rather successful. Replacement of the endogenous dH_4_MPT-dependent formaldehyde oxidation pathway with a glutathione-dependent pathway from *Paracoccus denitrificans* restored growth on methanol (59) but a series of several beneficial mutations acquired via experimental evolution was required to achieve growth at 80% that of wild-type (86–88). Introduction of dicholormethane (89) or chloromethane (90) metabolism was modestly successful, but a single beneficial mutation brought dichloromethane growth to the same as seen for natural *Methylobacterium* strains that use that substrate (91). Introduction of the large *mau* operon encoding the genes for methylamine dehydrogenase and its assembly was sufficient, without any further mutations, to achieve growth at the levels of natural *Methylobacterium* strains with that pathway (92). Experimental evolution of *M. extorquens* with pLC291-*pca-van* could reveal just how much better this strain could become as a VA conversion platform.

Finally, this study can be thought of as being analogous to the multiple natural instances of horizontal gene transfer of the *van*-*pca* pathway into *Methylobacterium* strains. Besides the *M. extorquens* AMS-5 strain noted before for growth on aromatics (44) a recently posted manuscript found several *M. extorquens* strains isolated from soybeans that also possess the ability to grow on VA (46). Our data suggest that methylotrophs may be particularly well pre-adapted to become consumers of methoxylated aromatics due to a propensity to handle formaldehyde toxicity and consume TCA cycle intermediates. This modularity provides a great advantage in terms of being able to couple the engineered aromatics metabolism studied here with other work developing *M. extorquens* PA1 for production of various products (55). Furthermore, given that the presence of these pathways simply provides resistance to the presence of VA and PCA, there may even be a selective advantage to protection that goes above and beyond the ability to grow on these substrates that are common in the plant and soil environments.

## Supplemental Figure(s)

**Figure S1:**
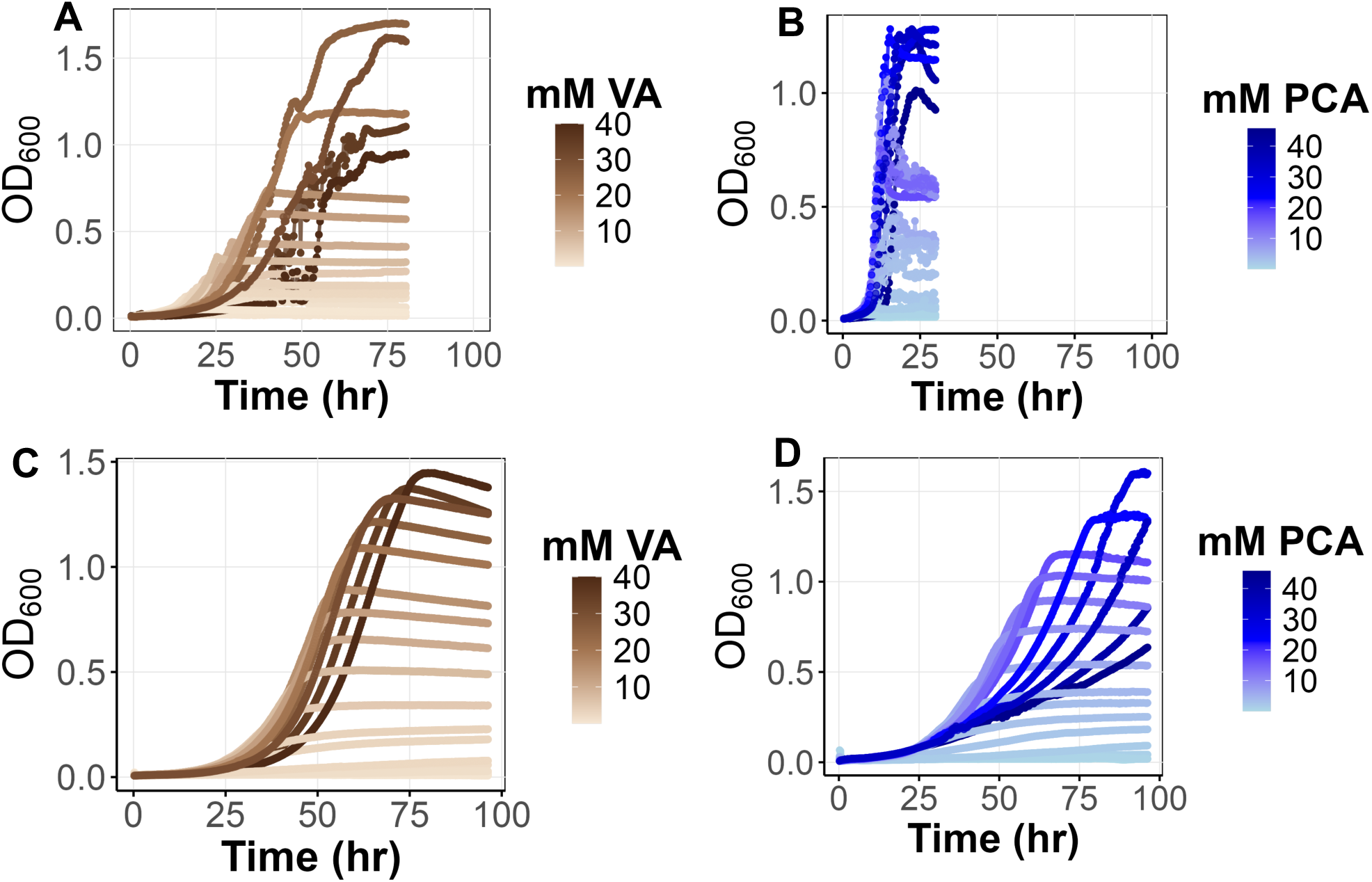
Comparative growth of *M. nodulans* and *M. extorquens* on vanillate VA and PCA. Growth curves of *M. nodulans* (A, B) and *M. extorquens* (C, D) with increasing concentrations of VA (A, C, red-yellow gradient) and PCA (B, D blue gradient).

**Figure S2:**
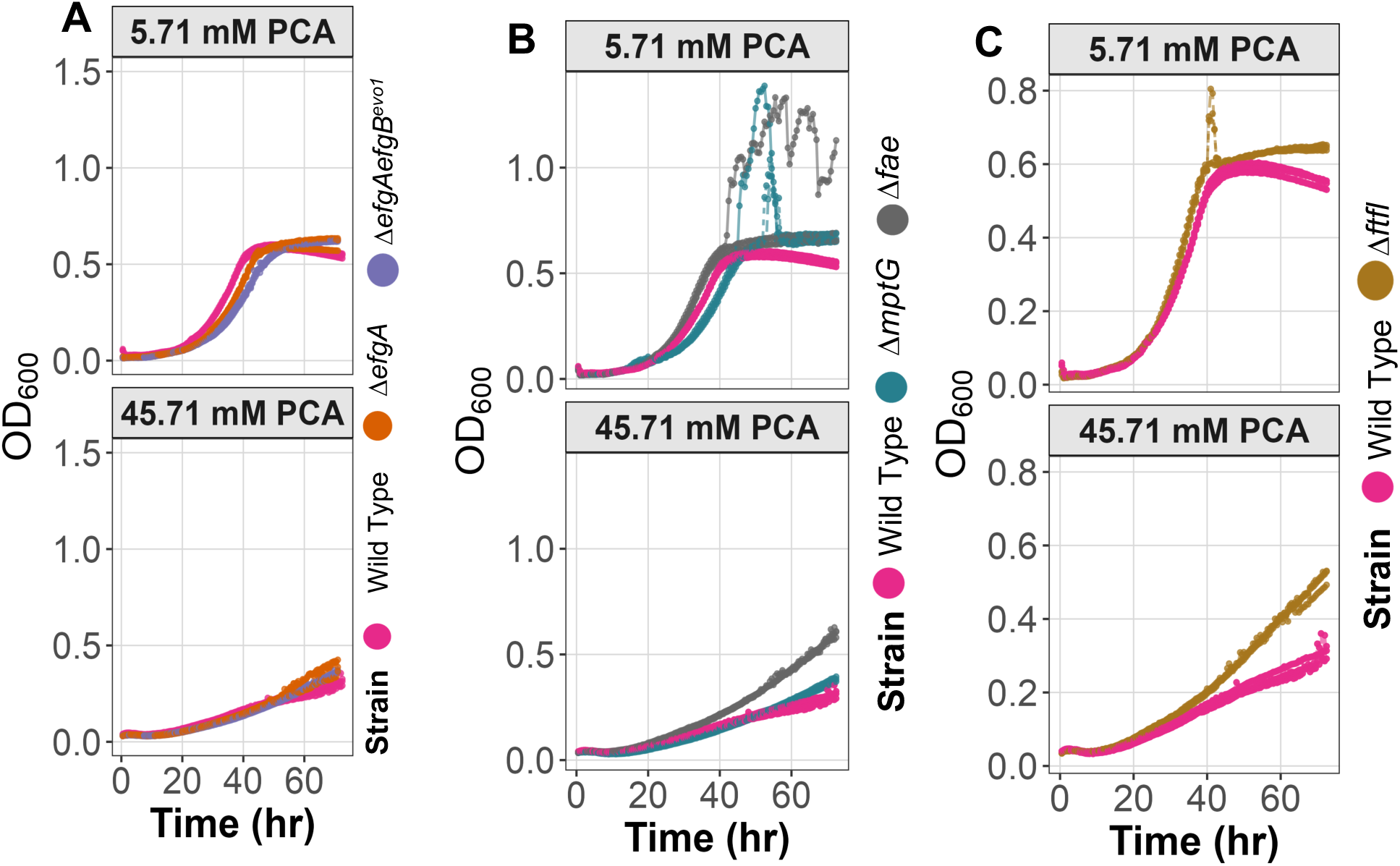
Growth curves of formaldehyde evolved alleles (A), oxidation mutants (B) and assimilation mutants (C) on PCA. None of the VA defects or advantages noted for these strains carried over to PCA, indicating their specificity to VA. Some strains exhibited new phenotypes on PCA, such as an advantage for the Δi*fae* strain, for reasons which we do not yet understand.

